# Analysis of vascular architecture and parenchymal damage generated by reduced blood perfusion in decellularized porcine kidneys using a gray level co-occurrence matrix

**DOI:** 10.1101/2021.10.18.464795

**Authors:** Igor V. Pantić, Adeeba Sharkeel, Georg A. Petroianu, Peter R. Corridon

## Abstract

There is no cure for kidney failure, but a bioartificial kidney may help address this global problem. Decellularization provides a promising platform to generate transplantable organs. However, maintaining a viable vasculature is a significant challenge to this technology. Even though angiography offers a valuable way to assess scaffold structure/function, subtle changes are overlooked by specialists. In recent years, innovative image analysis methods in radiology have been suggested to detect and identify subtle changes in tissue architecture. The aim of our research was to apply one of these methods based on a gray level co-occurrence matrix (GLCM) computational algorithm in the analysis of vascular architecture and parenchymal damage generated by hypoperfusion in decellularized porcine. Perfusion decellularization of the whole porcine kidneys was performed using previously established protocols. We analyzed and compared angiograms of kidneys subjected to pathophysiological arterial perfusion of whole blood. For regions of interest (ROIs) covering kidney medulla and the main elements of the vascular network, five major GLCM features were calculated: angular second moment as an indicator of textural uniformity, inverse difference moment as an indicator of textural homogeneity, GLCM contrast, GLCM correlation, and sum variance of the co-occurrence matrix. In addition to GLCM, we also performed discrete wavelet transform analysis of angiogram ROIs by calculating the respective wavelet coefficient energies using high and low-pass filtering. We report statistically significant changes in GLCM and wavelet features, including the reduction of the angular second moment and inverse difference moment, indicating a substantial rise in angiogram textural heterogeneity. Our findings suggest that the GLCM method can be successfully used as an addition to conventional fluoroscopic angiography analyses of micro/macrovascular integrity following in vitro blood perfusion to investigate scaffold integrity. This approach is the first step toward developing an automated network that can detect changes in the decellularized vasculature.

## 1 Introduction

The incidence of kidney failure, otherwise known as end-stage renal disease (ESRD) is rising globally (Arikan et al., 2021; Kari et al., 2021). Unfortunately, there is no cure for this condition, which can develop form the progression of acute and chronic injuries (Hsu and Hsu, 2016; Kolb et al., 2018; Corridon et al., 2021). Currently, transplantation is the best option to treat ESRD. Nevertheless, very few patients receive a timely transplant due to the complexity of the procedure, lack of donors, low viability of organs, and prevailing immunological incompatibilities (Saidi and Hejazii Kenari, 2014; Job and Antony, 2018; Wu et al., 2021). As a result, there is a definite need for alternatives to address this worldwide problem. Whole organ bioengineering has been proposed as one such alternative. Major advancements in this field have been developed using three-dimensional bioprinting, advanced stem cell technologies, and organ decellularization. Among these advancements, decellularization techniques currently hold the most promise for creating a bioartificial kidney (Sohn et al., 2020).

Decellularization is a unique alternative to porous scaffold fabrication systems, additive manufacturing procedures, and hydrogels, as it provides the necessary physical and biochemical environments to facilitate cell and tissue growth. This technology has garnered much attention within the past decade, as acellular scaffolds have been generated using bovine, equine, leporine, murine, and porcine models. However, substantial compromises to the scaffold architecture, observed under physiological conditions, inhibit their long-term viability and clinical utility (Zambon et al., 2018; Corridon, 2021). Thus, further research is needed to overcome problems related to vascularization and help realize the promise of a bioartificial kidney (Feng et al., 2020). Using this assertion, it is necessary to devise methods to better evaluate vascular patency in post-transplantation settings. Imaging modalities like X-ray/computed tomography, magnetic resonance imaging, ultrasonography and positron emission tomography have been applied to investigate the decellularized vascular architecture (Huling et al., 2016). These techniques provide useful information on the scaffold structure and function, as well as insight on the deformation that can arise after transplantation (Corridon, 2021). Yet, the low spatial resolution, artifacts, and unwanted morphological alterations have always proved to be challenging to detect subtle defects (Mostaco-Guidolin et al., 2013). Such challenges have paved the way for radiomic approaches that can extract features far beyond the capability of the human eye or brain to appreciate (Neri et al., 2019).

Computer-automated mathematical image analysis methods have emerged to give potentially wide applications in radiology. In recent years, many innovative techniques, and algorithms have been proposed and tested, often with limited success regarding their potential for integration in current diagnostic and research protocols. Future developments in information technology ensure that many of these techniques will significantly improve diagnostic and prognostic accuracies in X-ray computed tomography, fluoroscopy, and angiography (Cao et al., 2019; Kolossvary et al., 2021; V et al., 2021). Computational methods that use statistical analyses in evaluating image texture are potentially instrumental in X-ray imaging since they may enable fast, objective, and accurate detection of subtle changes in tissue architecture that are occasionally hard to notice during the conventional assessment. One such method is based on the gray level co-occurrence matrix (GLCM) algorithm, which has attracted much attention in computational medicine. The technique uses second-order statistics to determine indicators that reflect image features such as textural homogeneity, uniformity, and level of disorder. Previously, some of these indicators, such as angular second moment and inverse difference moment, have proven to be sensitive in assessing data obtained as the result of various X-ray digital image transformations (Chen et al., 2021; Kolossvary et al., 2021; Shankar et al., 2021).

In angiography, GLCM was successfully used as an addition to volumetric and radiomic metrics and image reconstruction of coronary lesions (Kolossvary et al., 2019). Also, some authors have previously demonstrated the potential of this method to evaluate endoleaks in aneurysmatic thrombus CT images of abdominal aorta (Garcia et al., 2012). Endovascular aortic aneurysm repair evolution might also be indirectly assessed with the help of GLCM and other textural algorithms (Garcia et al., 2014). Finally, in some experimental animal models, this form of textural analysis may be used to research pulmonary parenchymatous changes associated with pulmonary thromboembolism (Marschner et al., 2017). To the best of our knowledge, no such applications of GLCM have been used in evaluating kidney vascular architecture.

The aim of our work was to apply a gray level co-occurrence matrix GLCM computational algorithm to collectively assess vascular architecture and parenchymal damage generated from hypoperfusion in decellularized porcine kidneys using fluoroscopic angiography. We present evidence that GLCM may be highly applicable in the evaluation of normal and pathological kidney angiograms indicating its potential for inclusion in contemporary research practices in this area of radiology. Also, this is the first study to quantify textural changes in vascular architecture in decellularized kidney scaffolds, serving as the useful basis for future research on this organ model. Overall, this approach is the initial step toward developing an automated network that can detect changes in the decellularized vasculature.

## 2 Materials & Methods

### 2.1 Experimental animals

Adult Yorkshire pigs were euthanized, and whole kidneys were harvested under the guidelines provided by the Institutional Animal Care and Use Committee (IACUC) at the School of Medicine, Wake Forest University. All experimental protocols followed the ethical guidelines and regulations approved by Wake Forest University and the Animal Research Oversight Committee (AROC) at Khalifa University of Science and Technology. Moreover, all methods were performed in accordance with the Animal Research: Reporting of *In Vivo* Experiments (ARRIVE) guidelines.

### 2.2 Porcine kidney perfusion decellularization and sterilization

Whole porcine kidneys were extracted with intact renal arteries, veins, and ureters. The kidneys were then decellularized and sterilized using previously established protocols (Sullivan et al., 2012; Zambon et al., 2018; Corridon, 2021). Briefly, triton X-100, SDS, and phosphate-buffered saline (PBS) were slowly infused into cannulated renal arteries at a constant rate of 5 ml/min. Initially, 1% Triton X-100 was perfused through the renal artery for 36 h followed by 0.5% SDS dissolved in PBS for another 36 h. Finally, to remove the residual traces of detergents and cellular components, PBS was perfused through the kidneys for 72 h. The decellularized scaffolds were then submerged in PBS and sterilized with 10.0 kGy gamma irradiation.

### 2.3 Blood perfusion studies

Blood perfusion studies were carried out as previously reported (Corridon, 2021). Prior to perfusion, the bioreactor components, namely, suction pump heads (Ismatec, Cole-Palmer, Wertheim, Germany), standard pump tubing female and male luer x1/8ʺ hose barb adapters, barbed fittings, reducing connectors, three-way stop cocks, and Kynar adapters (Cole-Palmer, Vernon Hills, IL, USA) were sterilized using a 60Co Gamma Ray Irradiator. While the bioreactor tubing, chambers, and 2000 ml round wide mouth media storage bottles with screw caps assemblies (Sigma-Aldrich, St. Louis, MO, USA) were autoclaved.

Once sterilized, the bioreactor systems were assembled within a biosafety cabinet as described earlier (Corridon, 2021). Concisely, the chamber was assembled in a way that ensured the two outer blood flow lines were attached on either side of the suction pump head. This aided arterial outflow from the Ismatec MCP-Z Process or MCP-Z Standard programmable dispensing pump (Cole-Palmer, Vernon Hills, IL, USA) into the chamber’s arterial line while the venous returns to the pump via the venous line. The scaffold was suspended in a reservoir of roughly 500 ml of heparinized pig whole blood in the bioreactor chamber (BioIVT, Westbury, NY, USA). The renal artery was attached to the arterial line inside the chamber. In comparison, the renal vein cannula remained detached to allow venous outflow from the scaffold into the reservoir. The venous line was freely suspended into the reservoir to support unreplenished and unfiltered blood recirculation through the dispensing pumps. The entire assembled bioreactor system was then placed in a cell culture incubator, and scaffolds were subjected to continuous hypoperfusion (at a rate < 500 ml/min) for 24 hrs. At the 24-h time point, perfusion was ceased, and the scaffolds were removed from the chambers and placed in 60 x 15 mm sterilized polystyrene Petri dishes (Sigma-Aldrich, St. Louis, MO, USA) for fluoroscopic angiography.

### 2.4 Fluoroscopic angiography

Native and decellularized kidneys were first infused with 100 ml PBS via the renal artery. The contrast agent was infusion of Iothalamate meglumine contrast agent (60% Angio-Conray, Mallinckrodt Inc., St Louis, MO, USA). Once a sturdy flow of exiting contrast agent was achieved, the renal vein, renal artery, and ureter were occluded to prevent the contrast agent from leaking out of the organ. Angiograms were collected at ambient temperature in a sterilized suite with a Siemens C-arm Fluoroscope (Siemens AG, Munich, Germany).

### 2.5 GLCM analysis

We performed GLCM analysis of selected regions of interest in angiograms using Mazda computational platform. This software was created by Michal Strzelecki and Piotr Szczypinski of the Institute of Electronics, Technical University of Lodz (Szczypinski et al., 2007; Szczypinski et al., 2009; Strzelecki et al., 2013), Poland as a part of COST B21 European project “Physiological modelling of MR Image formation”, and COST B11 AQ6 European project “Quantitative Analysis of Magnetic Resonance Image Texture” (1998–2002). The software, originally made using C++ and Delphi© programming languages can accurately calculate GLCM features on multiple regions of interest (ROIs) of high-resolution BMP images making it an ideal candidate for textural analysis of angiograms.

In our angiograms in BMP format (bit depth equaled 24), we formed ROIs covering kidney medulla and the main elements of the vascular network, with the area of approximately 80000 resolution units (width of 200 and height of 400 resolution units) as shown in **Figure 2B**. For each ROI, five major GLCM features were calculated: angular second moment, inverse difference moment, GLCLM contrast, GLCM Correlation and Sum variance of the co-occurrence matrix. GLCM method assigns values to resolution units depending on their gray intensity, after which a series of complex second-order statistical calculations are performed on resolution unit pairs considering their distance and orientation. Values of individual GLCM features depend on the distribution patterns of the gray intensity pairs and the numerical organization of the resulting co-occurrence matrix.

**FIGURE 1.**
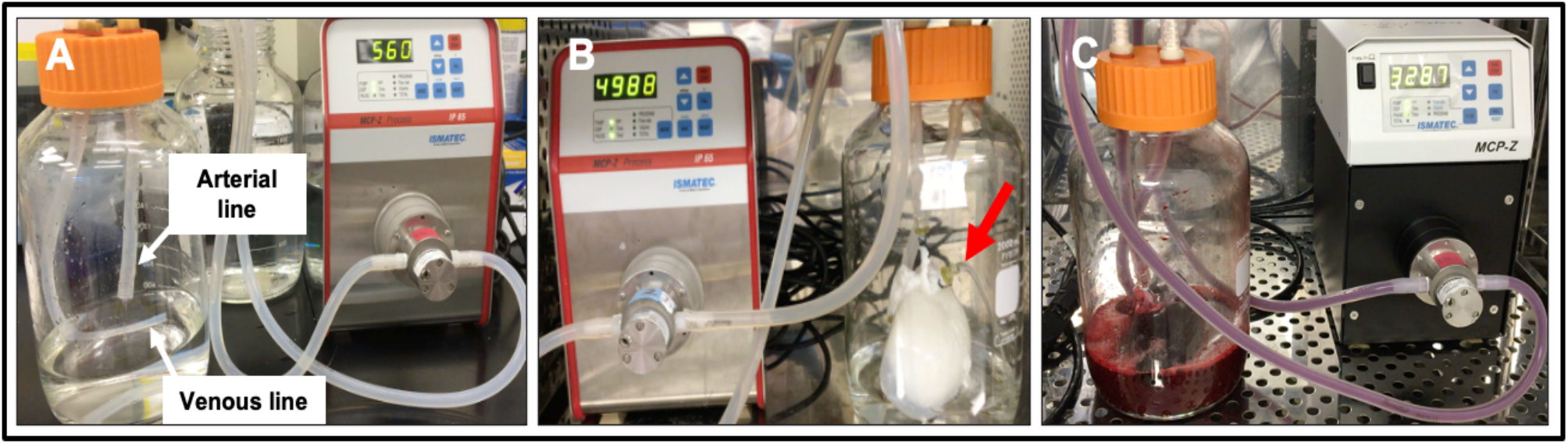
Photographs of the bioreactor used to perfuse decellularized scaffolds with whole blood (A) Image outlining the arterial line (which was then attached to the cannulated renal artery) and venous lines (which was left open to act as a venous reservoir to facilitate fluid recirculation) before the addition of the scaffold. (B) Image of an acellular kidney perfused with PBS illustrates how the scaffold recirculated fluid that emanated from its renal vein (red arrow) and open-ended venous line. (C) Image of a scaffold being perfused with whole pig blood.

**FIGURE 2.**
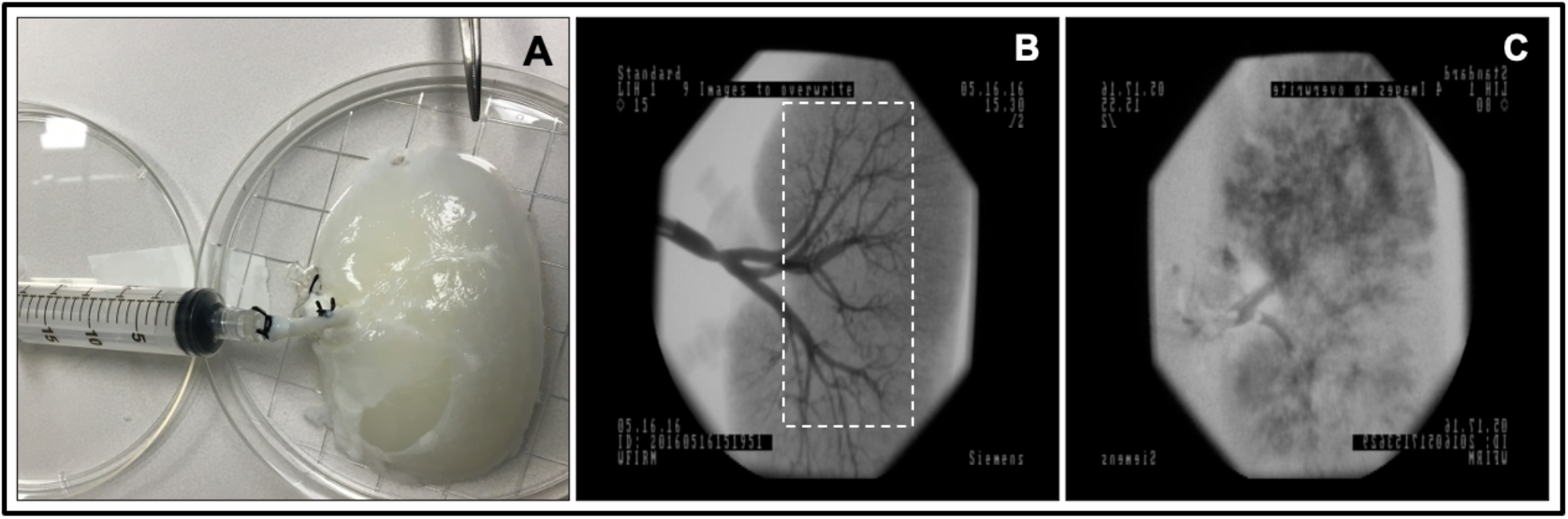
Fluoroscopic angiography. (A) Photograph of a decellularized scaffold that was set to be infused with contrast agent. (B) An angiogram of the scaffold before it was perfused with blood displaying the decellularized vascular network and region of interest (ROI), dashed rectangular region, covering kidney medulla and the main elements of this network. (C) An angiogram of the scaffold after 24 h of hypoperfusion (arterial infusion rate 20 ml/min).

In GLCM analysis, angular second moment (ASM) represents the level of textural uniformity in two-dimensional signal. It can be calculated as:

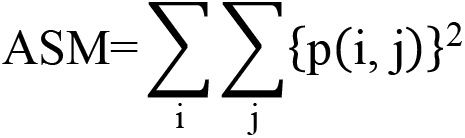

In this formula, p(i,j) is the (i,j)th entry of the gray-level co-occurrence matrix, after the normalization. In this work, angular second moment was in essence a tool for quantification of textural orderliness of the angiogram ROIs.

A relatively similar feature to angular second moment that can also be calculated during GLCM analysis is inverse difference moment. Inverse difference moment (Maidman et al.) is often used to quantify the level of textural smoothness, sometimes also referred to as “homogeneity”. It can be calculated as:

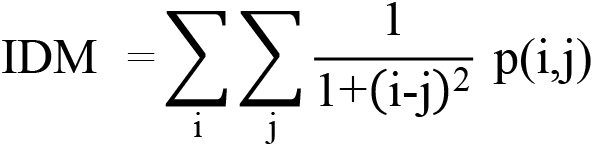

Some textural features take into account the mean (μ) and the standard deviation (σ) of normalized GLCM rows (i.e., *x* or *y*). Such is the GLCM Correlation parameter which is determined as:

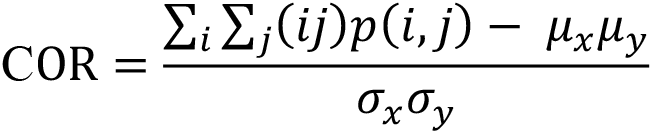

Textural sum variance feature is also a useful measure that can indirectly measure the level of dispersion around the mean of the matrix:

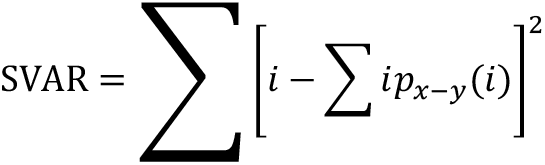

Finally, in our study, we also quantified the textural contrast feature as:

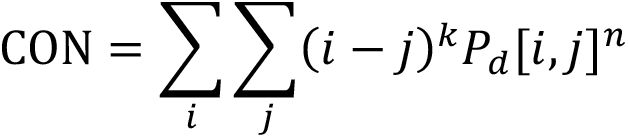

Textual contrast was used to quantify the difference between the neighbouring resolution units considering their respective gray intensities. For details on GLCM algorithm and the calculation of features, the reader is referred to previous works that deal on the application of this method in medical and other sciences (Haralick et al., 1973; Santos et al., 2015; Topalovic et al., 2021).

### 2.6 Discrete wavelet transform features

Discrete wavelet transform (DWT) analysis of angiogram ROIs was performed as an addition to calculation of GLCM features. The DWT algorithm in Mazda software includes linear transformation of data vectors to numerical vectors taking into account their lengths (in case of data vectors, the length of an integer power of two). The analysis is performed separately on rows and columns of data with the application of high (H) and low-pass (L) filtering (Kociolek et al., 2001). The final output of DWT includes energies (En) of wavelet coefficients (d) in different subbands (for a respective subband location x and y) at different scales for a ROI resolution unit number (n):

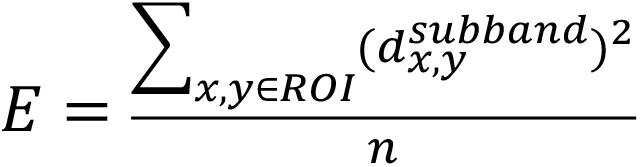

Previous research on application of DWT in microscopy has indicated that textural heterogeneity may influence the values of coefficient energies in subbands. In this work, we focused on the quantification of 3 such energies depending on the use of high (H) and low-pass (L) filtering: EnLH, EnHL and EnHH. Additional details on DWT algorithm can be found in previous publications (Kociolek et al., 2001; Paunovic et al., 2021a).

## 3 Results

### 3.1 Scaffold Perfusion Analyzed using Fluoroscopic Angiography and Venous Outflow

Fluoroscopic angiography showed that the vascular network was well-preserved post decellularization (**Figure 2B**). Angiograms taken from decellularized kidneys post-perfusion with unreplenished and unfiltered blood for 24 h revealed significant alterations in the decellularized vascular architecture and parenchyma (**Figure 2C**). The standard arterial branching patterns were noticeably disrupted by the end of perfusion, making it difficult to detect and differentiate the different branches of the arterial tree. Substantial levels of contrast agent extravasation were also observed throughout the cortical and medulla regions highlighting deleterious modifications to the decellularized renal parenchyma. Moreover, the hypoperfused acellular kidneys were unable to perfuse blood throughout their vascular networks and showed notable signs of thrombosis and cessation of venous outflow.

### 3.2 GLCM Analysis

The average angular second moment of the ROIs was 0.030 ± 0.018 for decellularized kidneys before perfusion (pre-perfusion) and 0.053 ± 0.022 for after perfusion (post post-perfusion) angiograms (**Figure 3A**). Statistically highly significant difference was observed (p < 0.01). This result implied a substantial reduction of textural uniformity in post-perfusion vascular architecture. Similar reduction was observed with the mean values of inverse difference moment (0.781 ± 0.046 in post-perfusion compared to 0.805 ± 0.026 in controls) (**Figure 3B**). The difference was significant (p < 0.05) which implied that the textural homogeneity of ROIs decreased.

**FIGURE 3.**
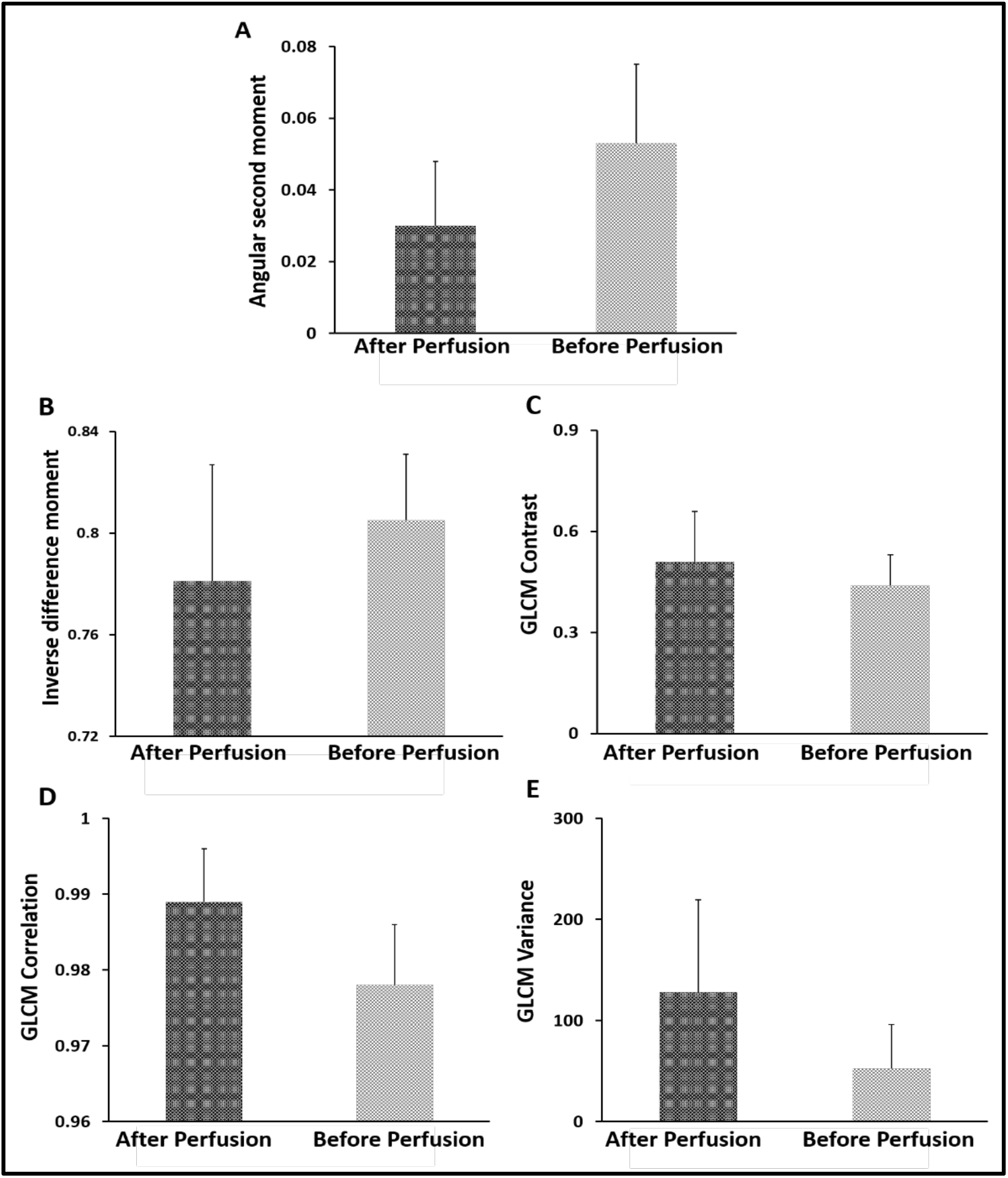
GLCM analysis. (A) Average angular second moment. (B) Inverse difference momentum. (C) GLCM contrast. (D) GLCM correlation. (E) GLCM variance.

On the other hand, there was a substantial rise in the average values of GLCM Contrast (**Figure 3C**), GLCM Correlation feature (**Figure 3D**) and Sum variance (**Figure 3E**). The largest increase was observed in Sum variance (127.99 ± 91.53 in post-perfusion versus 52.48 ± 43.36 in pre-perfusion angiograms, p < 0.01) followed by the Correlation (0.989 ± 0.007 versus 0.978 ± 0.008, p < 0.01), and lastly GLCM Contrast (0.51 ± 0.15 versus 0.44 ± 0.09, p < 0.05). This is in line with the results of the values of angular second moment and inverse difference moment, and imply the rise of the overall textural heterogeneity of vascular architecture.

We also observed some changes in wavelet coefficient energies of the ROIs, however these changes were not as drastic as the ones exhibited by GLCM features. The average value of EnLH rose from 1.17 ± 0.45 in controls to 1.54 ± 0.70 in post-perfusion angiograms (p < 0.05). Similar rise and level of significance was observed for EnHL means (1.07 ± 0.36 compared to 1.37 ± 0.53 in controls, p < 0.05). Regarding EnHH, the average value in pre-perfusion angiograms was 0.08 ± 0.01 and in post-perfusion angiograms it rose to 0.09 ± 0.02 (p > 0.05).

Significant correlations were detected between GLCM and DWT parameters in both groups of angiograms. For example, statistically highly significant negative correlation (p < 0.01) existed between the DWT EnHL feature and the values of inverse difference moment. Similar statistically highly significant negative correlation (p < 0.01) was observed between EnHH feature and the values of angular second moment. These associations are expected, they are partly the result of the methodological similarities during the implementation of GLCM and DWT algorithms, and confirm the validity of the obtained dataset.

## 4 Discussion

Time and again, emphasis has been given to the importance of maintaining the integrity of vascular networks in decellularized organs for transplantation. In this study, we applied textural analysis algorithms based on gray level co-occurrence matrix and discrete wavelet transform to investigate the pathologically changes in the vascular architecture of the decellularized kidneys subjected to conditions that mimic renal artery stenosis (Corridon, 2021). This form of stenosis is a well-recognized disorder that compromises transplantation and has been shown to denature the acellular vascular tracks. Such deformation also leads to aberrant changes in the decellularized parenchyma. The images obtained using fluoroscopic angiography showed that significant differences in both GLCM and DWT features could be detected using this approach. Our findings imply that textural analysis, as a set of contemporary and innovative computer-based methods, has a great potential to be used as an addition to the conventional angiographic evaluation of the renal vascular network.

Perhaps the most important finding was the observed significant change of inverse difference moment of angiogram ROIs. This GLCM feature indicates textural homogeneity and is often used to quantify smoothness in the distribution of resolution units in grayscale images. Previous research articles in digital micrographs have shown the potential value of inverse difference moment in detecting structural alterations that are not visible to a professional pathologist (Paunovic et al., 2021b). Along with angular second moment and GLCM contrast, this is probably also one of radiology’s most frequently calculated textural features.

Generally, in the past, the most frequent application of textural computational algorithms in radiology was to assess images obtained through nuclear magnetic resonance, computerized tomography, and other tomography techniques. Probably the most common approach is to compare the images of tissue lesions or other pathological changes in tissue architecture with controls. After that, one of the possibilities is to determine the sensitivity of individual GLCM features for lesion detection or to test the discriminatory power of the method regarding the separation of post- and pre-perfusion radiographs or parts of a radiograph (Chen et al., 2021). Another strategy would be to use GLCM features as prediction tools for disease prognosis (Huang et al., 2021), or to test their ability to determine boundaries of the lesion in the same radiograph. Finally, it may be possible to develop a scoring system that considers GLCM (and other) indicators of texture and test its sensitivity and specificity (Thuillier et al., 2021).

In angiography, the GLCM method is much less frequently applied, and so far, only a handful of studies have been published on this topic. These mainly include the use of GLCM features for assessment of low attenuation noncalcified (LANCP), noncalcified and calcified coronary plaques (Kolossvary et al., 2019; Kolossvary et al., 2021), or for computer-aided diagnosis-specific cases of endovascular aortic aneurysms (Garcia et al., 2014). In optical coherence tomography angiography, as demonstrated earlier, some GLCM indicators can also be applied to quantify choriocapillaris in healthy and diseased eyes (Khan et al., 2020). To the best of our knowledge, there hasn’t been a similar study trying to apply texture analysis for the assessment of vascular changes in kidney tissue. Therefore, our research is probably the first to demonstrate the applicability of these computational algorithms (GLCM and DWT techniques on an experimental model of decellularized kidney) in this rapidly developing area of radiology and also provides a potentially useful foundation for future research.

In the future, probably the most important application of both GLCM and DWT analyses will be to provide inputs for various artificial intelligence-based methods for image analysis in radiology. This application would include training and testing different machine learning models, some of which have already been suggested as suitable for GLCM data (Davidovic et al., 2021). The examples would be conventional decision tree algorithms such as CHAID (Chi-square Automatic Interaction Detector) or CART (classification and regression tree) or some more modern approaches such as random forests. Support vector machines, naive Bayes, linear discriminant analysis, and similarity learning are potential alternative strategies. The most considerable potential regarding the use of GLCM and DWT raw data may lie in designing various types of neural networks. This process includes simple concepts such as a multilayer perceptron or more complex ones such as recurrent and convolutional neural networks. Convolutional neural networks are a fascinating approach since they are already widely used in medicine and other disciplines of computer vision. Despite the promises that such a computer algorithm makes, many loopholes still need to be addressed that will require extensive quality assurance of these methods, including testing inter- and intra-observer reliability, for their effective application in the clinics.

As mentioned, the limitations of our study include the relatively small sample, which is not sufficient for the implementation of the more complex approaches such as machine learning or the creation of other artificial intelligence-based models. Also, another important aspect to consider is that the results of GLCM and DWT generally depend on various factors associated with image creation. Brightness, contrast, hue, saturation, and many other image parameters which can vary in angiograms can substantially impact GLCM features such as angular second moment or inversed difference moment. Finally, to our knowledge, the results of the textural analysis are not always the same across different computational platforms. Such variations can arise from the fact that existing software algorithms may use images in different formats (8-bit, 16-bit, BMP, JPG, etc.) or because of many other technical issues and solutions the developers tried to include into the programming code. All of these issues may in the future hinder the potential of successful integration of textural analysis methods in contemporary diagnostic protocols. For this to happen, extensive quality assurance of the processes, including testing inter- and intra-observer reliability, will have to be performed.

## 5 Conclusion

Our results designate that certain discrete changes in vascular architecture and renal parenchyma in the decellularized kidney can be successfully detected using contemporary innovative computational algorithms for texture analysis, thereby overcoming the limitations of conventional imaging modalities. We report statistically significant changes in GLCM and wavelet features, including reducing angular second moment and inverse difference moment, indicating a substantial rise in angiogram textural heterogeneity in pathological conditions. Our findings suggest that the GLCM method may be used as an addition to the conventional fluoroscopic angiography analysis of micro‑/macrovascular integrity for a more accurate diagnosis. To the best of our knowledge, this is the first study to use GLCM and DWT based approach in decellularized kidney experimental model, augmenting appropriate evaluation of the decellularized kidneys vasculature, to accomplish lasting vascular patency post-transplantation, thereby giving hope to impede a looming epidemic of morbidity or mortality due to kidney diseases. This approach is the first step toward developing an automated network that can detect debilitating changes in the decellularized vasculature and supporting tissue network.

## 7 Author Contributions

P.R.C. and I.V.P, conceived and designed project. P.R.C. performed all experiments, analyzed the associated data, interpreted results of experiments. I.V.P. performed all computation analyses. I.V.P., A.S., G.P., and P.R.C drafted, edited, and approved final version of manuscript.

## 9 Acknowledgments

The author would like to acknowledge Dr. Zambon for help with decellularization. The authors would also like to thank Ms. Anousha Khan, Ms. Xinyu Wang, and Nnamdi Ugwuoke for reviewing the manuscript.

## 10 Data Availability Statement

The datasets generated for this study are available on request to the corresponding author.

